# The plasma biomarker soluble SIGLEC-1 is associated with the type I interferon transcriptional signature, ethnic background and renal disease in systemic lupus erythematosus

**DOI:** 10.1101/266965

**Authors:** João J. Oliveira, Sarah Karrar, Daniel B. Rainbow, Christopher L. Pinder, Pamela Clarke, Arcadio Rubio García, Osama Al-Assar, Keith Burling, Tim J. Vyse, Linda S. Wicker, John A. Todd, Ricardo C. Ferreira

## Abstract

The molecular heterogeneity of autoimmune and inflammatory diseases has been one of the main obstacles to the development of safe and specific therapeutic options. Here we have evaluated the diagnostic and clinical value of a robust, inexpensive, immunoassay detecting the circulating soluble form of the monocyte-specific surface receptor sialic acid binding Ig-like lectin 1 (sSIGLEC-1).

We developed an immunoassay to measure sSIGLEC-1 in small volumes of plasma/serum from systemic lupus erythematosus (SLE) patients and healthy donors. Plasma concentrations of sSIGLEC-1 strongly correlated with expression of SIGLEC-1 on the surface of blood monocytes and with type I interferon (IFN)-regulated gene (IRG) expression in SLE patients. In addition, we identified marked ancestry-related differences in sSIGLEC-1 concentrations in SLE patients, with patients of non-European ancestry showing higher levels compared to patients of European ancestry. Higher sSIGLEC-1 concentrations were associated with lower serum complement component 3 and increased frequency of renal complications in European patients, but not with the SLEDAI clinical score.

Our sSIGLEC-1 immunoassay provides a specific and easily-assayed marker for monocyte-macrophage activation, and interferonopathy in SLE and other diseases. Further studies can extend its clinical associations and its potential use to stratify patients and as a secondary endpoint in trials.

## Introduction

The type I interferon (IFN) pathway was identified as a central feature of the autoimmune disease systemic lupus erythematosus (SLE) when IFN-α was first detected at high levels in patients’ sera (1). Since this initial observation, the development of SLE-like clinical manifestations in patients treated with IFN-α for different malignancies pointed to the involvement of IFN-α in the aetiology of the disease (2). Furthermore, naturally occurring anti-IFN-α antibodies in SLE patients have been shown to be associated with milder forms of the disease (3). The identification of a constitutive IFN transcriptional signature of hundreds of IFN-regulated genes (IRGs) in peripheral blood from a subset of SLE patients (4, 5) suggested that the IFN signature could be used as a clinical biomarker to stratify patients with autoimmune and inflammatory diseases in which type I IFNs are known to play a pathogenic role, referred to as the interferonopathies. Nevertheless, the precise link between the IFN signature and molecular subtypes of disease or with broader disease severity scores has been put into question (6). With the development of more sophisticated high-throughput genomic tools, it has become apparent that the IFN signature is a complex composite marker, which can be further stratified into several distinct signatures that are better predictors of disease subtype (7). Longitudinal analyses have revealed that the IFN signature is highly variable over time as a result of alterations in blood composition caused by therapy or progression of the disease (7, 8). This is particularly relevant to IFN-driven diseases, as IFN-α is known to significantly alter the relative distribution of immune cell types in blood, which can severely compromise the diagnostic potential of the whole blood IFN signature (8) and lead to the observed lack of correlation between the signature and disease activity over time (9, 10).

These findings have led to the investigation of other potential cell-type specific biomarkers that could be better predictors of disease severity or clinical subtypes. One such IFN-regulated marker that has shown promise for the stratification of SLE patients is sialic acid binding Ig-like lectin 1 (SIGLEC-1) (11–13). SIGLEC-1 is a cell-adhesion molecule involved in the initial contacts with sialylated pathogens and mediates phagocytosis and endocytosis of pathogens, thereby promoting efficient immune responses to limit infection (14). Unlike the canonical IFN transcriptional signature which is a composite of many genes expressed at different levels in different immune cells, SIGLEC-1 is expressed exclusively in cells of the myeloid lineage, namely tissue-resident macrophages and monocyte-derived dendritic cells (15, 16). In blood, expression of surface SIGLEC-1 is restricted to CD14^+^ monocytes, and has been previously reported to be increased in several other autoimmune diseases, including rheumatoid arthritis (17), systemic sclerosis (18) and primary biliary cirrhosis (19). In addition, a recent study has shown that increased SIGLEC-1 expression on the surface of monocytes was a predictor of congenital heart block during pregnancy in children from Ro/SS-A autoantibody-positive mothers (20).

These data support the use of SIGLEC-1 as a potential cell-type specific biomarker for the stratification of patients with an overt type I IFN response. However, assay of surface SIGLEC-1 requires intact cells and flow cytometry, features that are not conducive for the development of a high throughput, inexpensive biomarker assay, ideally detectable in plasma/serum. Here we show that a circulating form of SIGLEC-1 can be detected in serum/plasma, and developed a robust and inexpensive immunoassay to measure its concentration. Furthermore, we provide evidence that the concentration of soluble SIGLEC-1 (sSIGLEC-1) is associated with patient’s ancestry and with renal involvement in SLE patients. Therefore, sSIGLEC-1 is a new circulating plasma/serum biomarker of type I IFN activity in systemic autoimmune, inflammatory and infectious diseases that can be used accurately and conveniently in large numbers of samples, and could be used in clinical trials of drugs modulating the IFN signalling pathway for patient stratification and as a secondary endpoint.

## Results

### Soluble SIGLEC-1 assay development

To investigate whether we could detect SIGLEC-1 expression levels in peripheral blood, we developed an immunoassay based on time-resolved fluorescence (TRF) to measure the concentrations of the circulating form of the receptor, which we refer to as sSIGLEC-1. Although *SIGLEC1* is predicted to encode a soluble protein isoform, it has not been previously shown that such soluble protein can be detected in plasma/serum.

We found that sSIGLEC-1 was detected in circulation, with concentrations ranging from 1.29-276.1 ng/ml in plasma/serum samples from healthy donors and SLE patients. Technical variation of the assay was found to be very low, as assessed in two independent measurements of the same plasma sample from 23 healthy donors performed 308 days apart (median CV=4.8%, r^2^=0.91; Figure 1A), indicating minimal inter-assay variability. Similarly, biological sSIGLEC-1 levels were found to be very stable (CV = 11.8%, r^2^=0.67; Figure 1B) in 19 healthy donors bled at two separate visits (median time between visit, 378 days; range 239-519 days, with only two donors showing physiological differences (CV>20%) between visits, likely due to viral infections (21). Of note, one donor showed high concentrations of sSIGLEC-1 on the first visit (28.3 ng/ml), which were maintained after 343 days of the initial measurement (23.4 ng/ml; Figure 1B), suggesting that, in addition to viral and possibly non-viral infections, genetic factors regulate the sSIGLEC-1 levels.

**Figure 1.**
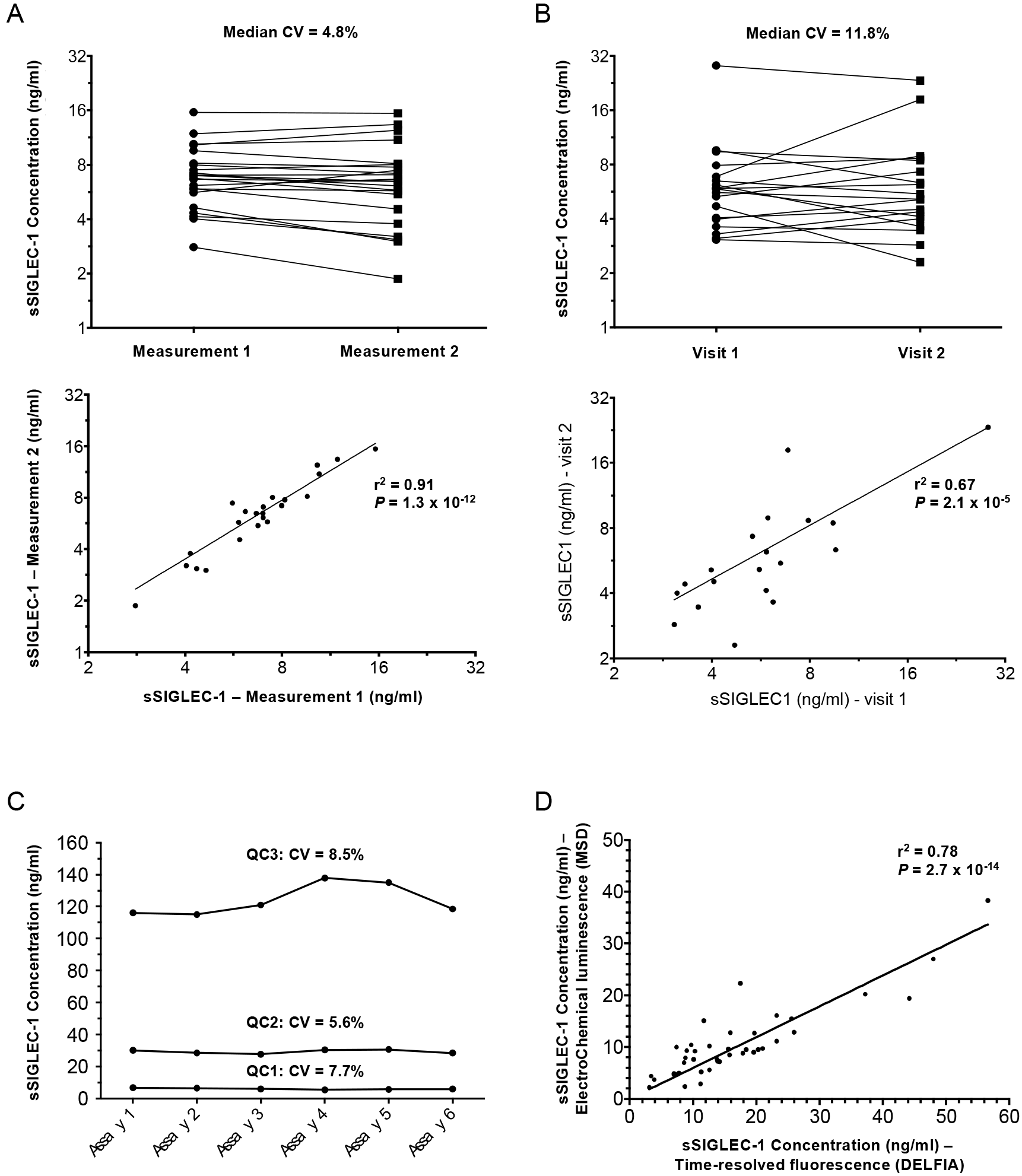
sSIGLEC-1 stability and assay performance. (A) Data shown depict the inter-assay (technical) variation of the time-resolved fluorescence (TRF) sSIGLEC-1 immunoassay. Data were obtained from the measurement of the same plasma sample from 23 healthy donors on two independent assays, performed 308 days apart. (B) Data shown depict the intra-individual (biological) variation of sSIGLEC-1 concentration between two visits of the same donor. Longitudinal physiological variation was assessed in 19 independent healthy donors using plasma samples collected at each separate visit. Median time between visits was 378 days (min = 239 days, max = 519 days). (C) Technical variation of the electrochemical luminescence (ECL) sSIGLEC-1 immunoassay using the MesoScale Discovery (MSD) platform. Stability of the assay was assessed in three QC pools of serum samples with increasing sSIGLEC-1 concentrations (QC1 = 6.0 ng/ml, QC2 = 29.2 ng/ml and QC3 = 123.9 ng/ml), which were measured in each assay over a two-day period. (D) Correlation between the TRF (DELFIA) and ECL (MSD)-based immunoassays. Measurements were performed in independent serum aliquots from a subset of 41 SLE patients from cohort 3. All correlations between measurements and *P* values were calculated by linear regression.

To expand the applicability of the sSIGLEC-1 immunoassay, we also developed an electrochemical luminescence-based (ECL) assay on the Meso Scale Discovery (MSD) platform, using the same detection antibodies and experimental protocol. The assay working range was found to be 0.5-1000 ng/ml, with an approximate 92% recovery of recombinant SIGLEC-1 protein spiked into serum samples. Stability of the assay over time was assessed using three pools of serum samples with increasing sSIGLEC-1 concentrations that were measured over six different assays run over a two-day period. Assay stability was consistent with the TRF assay, with CVs =7 .7%, 5.6% and 8.5%, for the low, medium and high QC pools, respectively (Figure 1C). Furthermore, we found a very high concordance between the TRF and ECL assays (r^2^ = 0.78; Figure 1D), as assessed by measuring a subset of 41 SLE patients on both platforms, using two independent serum aliquots.

### Soluble SIGLEC-1 levels are correlated with the surface SIGLEC-1 expression on CD14^+^ monocytes

Currently, the surface expression of SIGLEC-1 on monocytes has been suggested as a sensitive cell-type specific biomarker for SLE in blood (13). To investigate the relationship between surface and soluble SIGLEC-1 levels, we immunophenotyped the expression of this protein on the surface of CD14^+^ monocytes in peripheral blood mononuclear cells (PBMCs) collected from a discovery cohort (cohort 1) of 34 SLE patients and 24 matched healthy donors (Figure 2A). Consistent with previous findings (12, 13), we found that the surface expression of SIGLEC-1 was increased in CD14^+^ monocytes from SLE patients compared to healthy controls (*P* = 1.4 × 10^−4^; Figure 2B). However, we found bimodal expression of the surface SIGLEC-1 among SLE patients, with ten subjects (10/34: 29%; Figure 2B) presenting low levels of protein expression, similar to the ranges observed in healthy donors, and the rest presenting much higher levels, rarely observed in healthy volunteers (Figure 2A, B).

**Figure 2.**
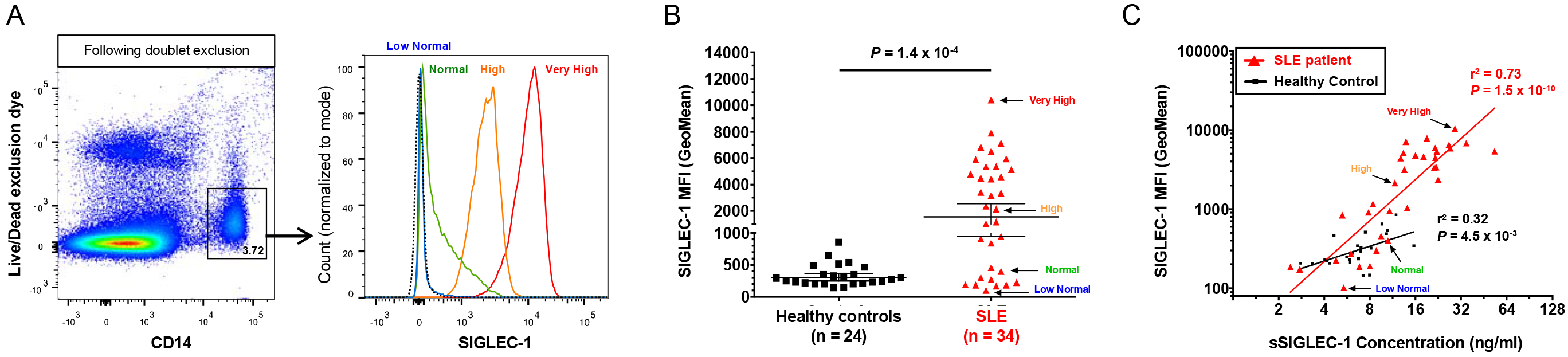
Soluble SIGLEC-1 is a surrogate for the surface expression of SIGLEC-1 on CD14^+^ monocytes. (A) Gating strategy for the delineation of CD14^+^ monocytes, following single-cell discrimination. Histograms depict the distribution of SIGLEC-1 expression on the surface of CD14^+^ monocytes obtained by flow cytometry in illustrative donors expressing: (i) Low Normal (blue); (ii) Normal (green); (iii) High (orange); or (iv) Very High (red) levels of SIGLEC-1. The dotted line represents the background expression of SIGLEC-1 in live lymphocytes, which are known not to express SIGLEC-1. (B) Scatter plot depict the frequency (geometric mean +/− 95% CI) of SIGLEC-1 expression on the surface of CD14^+^ monocytes in a discovery cohort (cohort 1) of healthy donors (N = 24; black squares), and SLE patients (N = 34; red triangles). *P* value was calculated using a two-tailed non-parametric Mann-Whitney test. (C) Correlation between the SIGLEC-1 Mean Fluorescence Intensity (MFI) on CD14^+^ monocytes obtained by flow cytometry and the corresponding the sSIGLEC-1 concentrations in the healthy control and SLE patient groups. *P* value was calculated by linear regression. The illustrative SIGLEC-1 Low Normal, Normal, High and Very High SLE patients shown inn (A) are highlighted in the plots in (B) and (C). SLE, systemic lupus erythematosus.

We found a strong correlation between the surface expression of SIGLEC-1 on monocytes and the concentration of sSIGLEC-1 in plasma samples from the same donors, particularly among SLE patients (r^2^ = 0.73, *P* = 7.9 × 10^−10^; Figure 2C). The distribution of sSIGLEC-1 levels recapitulated the same bimodal distribution of the surface SIGLEC-1 expression on SLE patients, ranging from ‘low normal’ physiological levels observed in healthy donors to the very high levels observed in a subset of patients (Figure 2C).

### Association of sSIGLEC-1 with the IFN transcriptional signature and SLE disease activity index

To assess if sSIGLEC-1 was associated with the IFN signature, we measured the transcription of 56 IRGs previously found to recapitulate the IFN signature (21). We found that sSIGLEC-1 levels were significantly correlated with the IFN transcriptional signature in PBMCs from SLE patients (r^2^ = 0.67, *P* = 2.9 × 10^−9^; Figure 3A). The correlation was also observed in healthy donors (r^2^ = 0.34, *P* = 2.8 × 10^−3^; Figure 3A), albeit to a lower extent. These findings were replicated in an independent cohort (cohort 2) of 41 SLE patients (r^2^ = 0.46, *P* = 1.2 × 10^−6^; Figure 3B), confirming that sSIGLEC-1 is a marker of the peripheral IFN signature. The lower correlation in the replication cohort is consistent with a lower overall disease severity – and concomitant lower sSIGLEC-1 levels – of the patients in the replication cohort, which were recruited outside their regular clinic visits.

**Figure 3.**
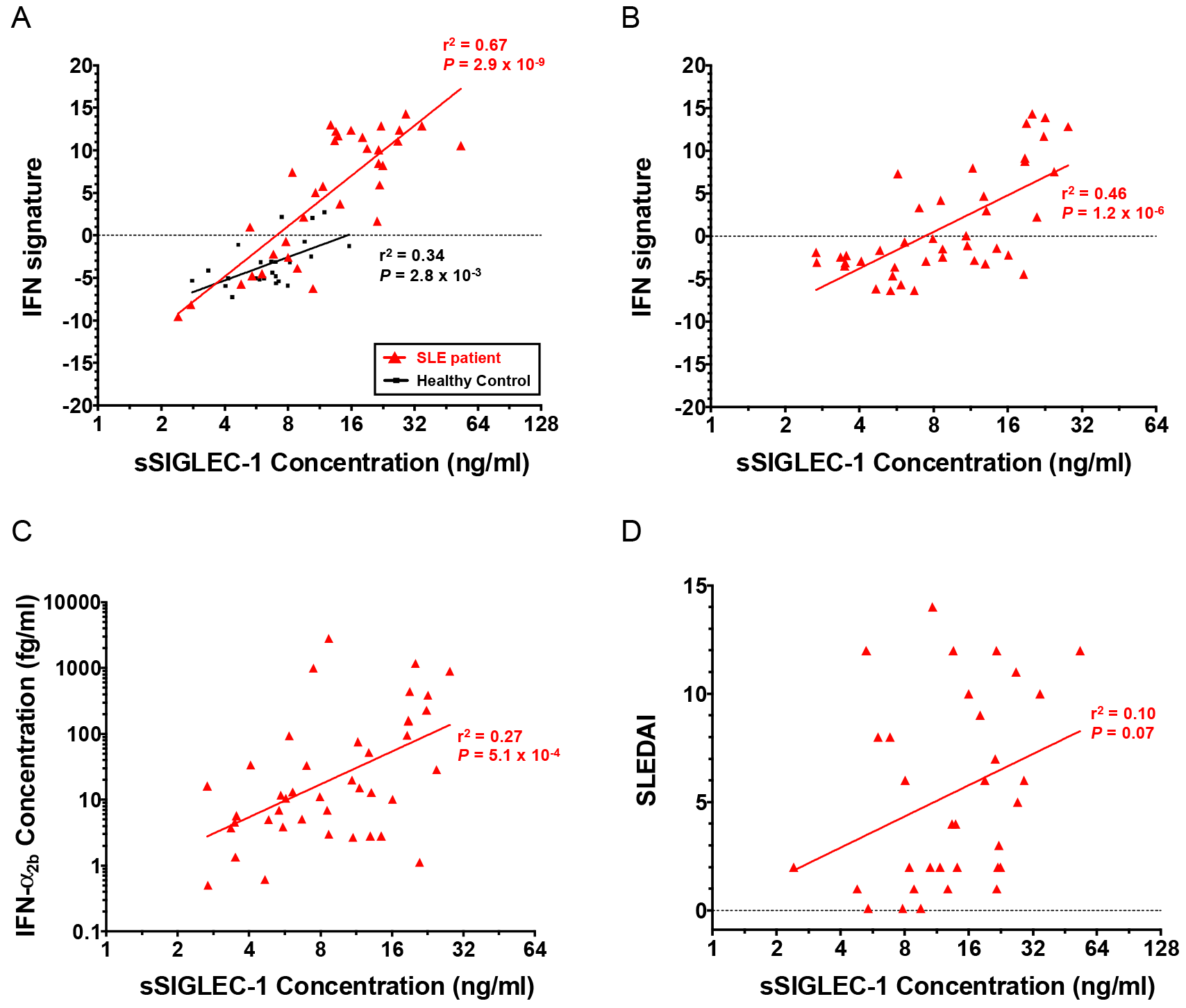
Comparison of sSIGLEC-1 with other markers of disease activity. (A,B) Correlation between the sSIGLEC-1 concentration and the canonical IFN transcriptional signature obtained by NanoString in RNA from the same donors. The IFN signature was measured in 24 healthy donors (depicted in black) and 34 SLE patients (depicted in red) from the discovery cohort (cohort 1) (A), and in the 41 SLE patients from the replication cohort (cohort 2) (B). (C) Correlation between the sSIGLEC-1 and the IFN-α_2b_ concentrations measured in plasma samples from the 41 SLE patients in cohort 2. (D) Data shown depicts the correlation between the sSIGLEC-1 concentration and the SLE disease activity score (SLEDAI) available from the 34 SLE patients from cohort 1. *P* values were calculated by linear regression. IFN, type I interferon.

Recently a single-molecule digital ELISA assay has been shown to detect IFN-α at femntomolar levels in circulation even from healthy individuals (22). Consistent with its potent biological activity, over a third of the SLE patients showed very low IFN-α_2b_ concentrations (< 10 fg/ml; Figure 3C), which were close to the reported limit of detection. In our hands the assay showed good reproducibility (CV = 4.1% between replicates), indicating that it is sensitive to detect even low concentrations of IFN-α_2b_. We found a significant correlation between the concentrations of IFN-α_2b_ in plasma and sSIGLEC-1 (r^2^ = 0.27, *P* = 5.1 × 10^−4^; Figure 3C) as well as the IFN signature (r^2^ = 0.34, *P* = 5.6 × 10^−5^; Supplemental Figure 1), although both were less pronounced than the observed correlation between sSIGLEC-1 and the IFN transcriptional signature.

In our sample of 34 SLE patients from the discovery cohort, sSIGLEC-1 concentrations and the SLE disease activity index (SLEDAI) score were not correlated (r^2^ = 0.10, *P* = 0.07; Figure 3D). This result is consistent with previous evidence showing a lack of association of other common serological disease markers, including various intra-nuclear autoantibodies, elevated B-cell activating factor of the tumour necrosis factor family (BAFF) levels and hypocomplementaemia, as well as the whole blood IFN signature, with disease activity scores such as SLEDAI (6).

### Increased sSIGLEC-1 concentration is associated with renal involvement

Having assessed that sSIGLEC-1 is an IFN-regulated marker that can be detected in circulation, we next investigated its potential as a clinical biomarker in SLE. Similarly to surface SIGLEC-1 expression, sSIGLEC-1 concentrations were markedly increased in SLE patients (10.4 ng/ml, 95% CI 8.8-12.2) compared to healthy controls (5.78 ng/ml, 95% CI 5.5-6.0, *P* = 9.6 × 10^−12^; Figure 4A) in the combined discovery and replication cohorts of 75 SLE patients and 504 healthy donors.

**Figure 4.**
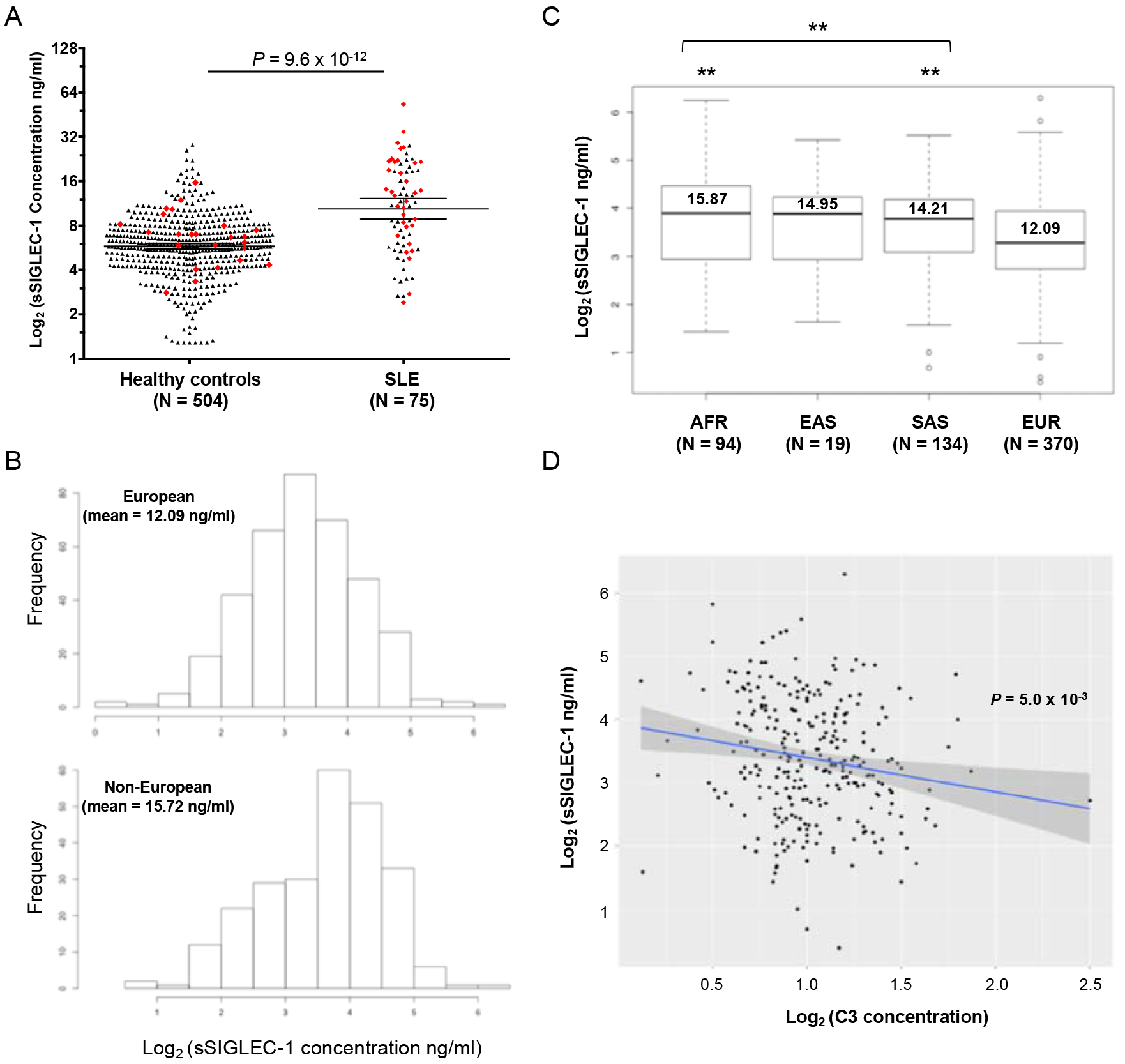
Association of sSIGLEC-1 with ancestry and serological markers of SLE. (A) Scatter plots depict the frequency (geometric mean +/− 95% CI) of the sSIGLEC-1 concentration in healthy donors (N = 504) and SLE patients (N = 75). The 24 healthy donors and 34 SLE patients from cohort 1 for which we have generated additional immunophenotyping and transcriptional data are depicted by red diamonds. *P* value was calculated using a two-tailed non-parametric Mann-Whitney test. (B) Histograms depict the distribution of sSIGLEC-1 concentrations in European and non-European SLE patients from cohort 3. (C) Box plots depict the distribution of sSIGLEC-1 concentration in the SLE patients stratified by ancestry group. Mean sSIGLEC-1 levels are indicated for each population. Twenty-three patients of additional minor ancestry groups (including Middle Eastern, Maori and Fiji) were included in the non-European population for the combined analysis. *P* values were calculated using Pearson’s chi-squared test comparing the concentration of sSIGLEC-1 of each individual of combined non-European populations to the patients of European ancestry. (D) Association between sSIGLEC-1 concentrations and serum complement 3 (C3) levels. *P* value was calculated by linear regression. \*\**P*<0.01; EUR, European; AFR, African/Afro-Caribbean; SAS, South East Asian; EAS, East Asian.

To assess the potential clinical application of sSIGLEC-1, we measured sSIGLEC-1 levels in serum from 656 SLE patients with available clinical information (cohort 3). We observed that the concentrations of sSIGLEC-1 were significantly higher in non-European patients (15.7 ng/ml, 95% CI 13.52-17.92) compared to those of European ancestry (12.1 ng/ml, 95% CI 11.25-12.94, *P* = 2.7 × 10^−3^; Figure 4B). Soluble SIGLEC-1 levels were similarly elevated among the different non-European populations (Figure 4C), and were therefore combined into a single group to increase statistical power. In agreement with their reported increased disease severity, non-European SLE patients showed increased frequency of renal and neurological complications (Table 1), thus supporting the hypothesis that increased underlying disease severity in non-European patients drives their increased levels of sSIGLEC-1.

**Table 1.**
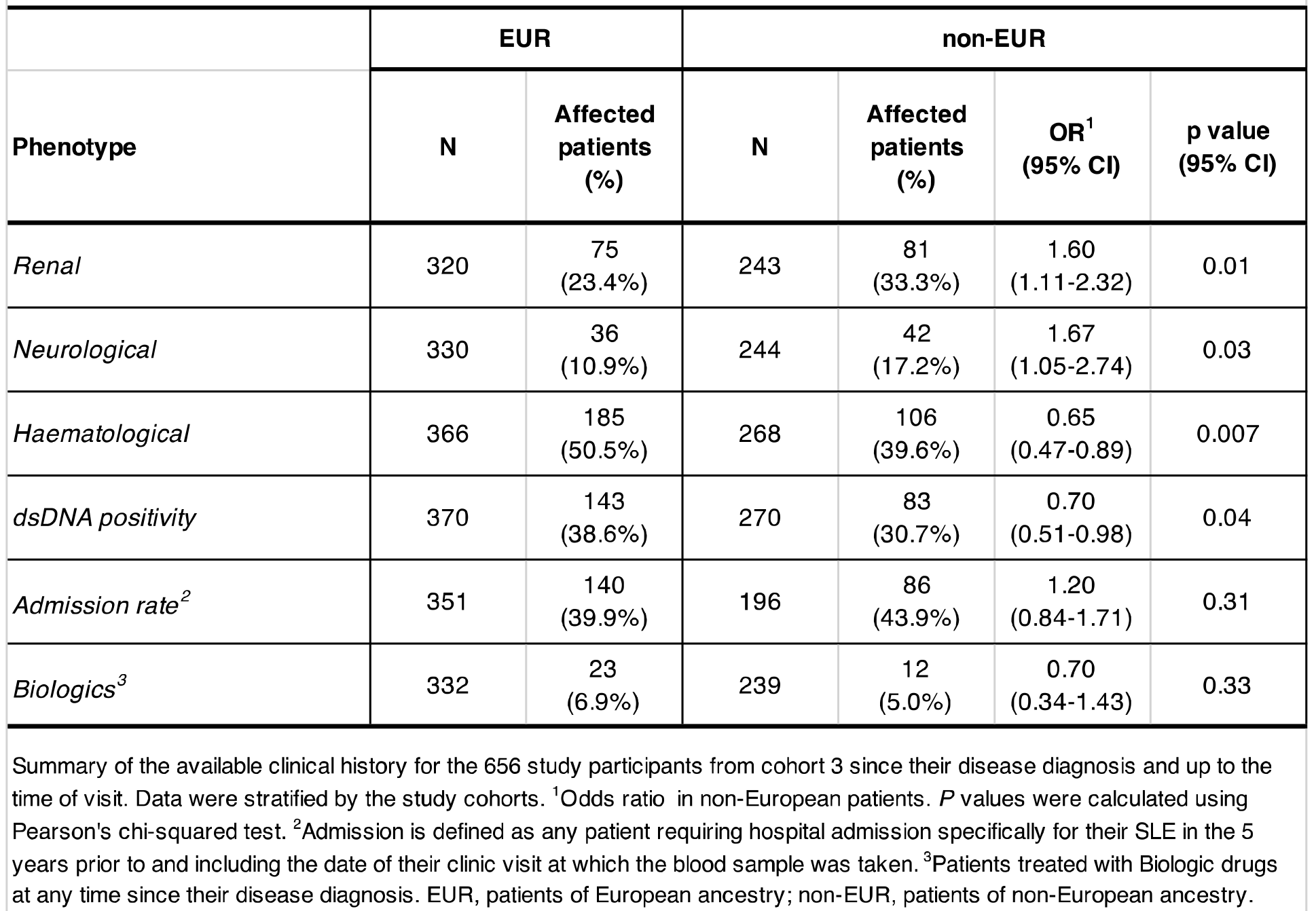
Summary of the history of clinical manifestation of the SLE cohort 3

|  | EUR |  | non-EUR |  |  |  |
| --- | --- | --- | --- | --- | --- | --- |
| Phenotype | N | Affected patients (%) | N | Affected patients (%) | OR<sup>1</sup> (95% CI) | p value (95% CI) |
| *Renal* | 320 | 75 (23.4%) | 243 | 81 (33.3%) | 1.60 (1.11-2.32) | 0.01 |
| *Neurological* | 330 | 36 (10.9%) | 244 | 42 (17.2%) | 1.67 (1.05-2.74) | 0.03 |
| *Haematological* | 366 | 185 (50.5%) | 268 | 106 (39.6%) | 0.65 (0.47-0.89) | 0.007 |
| *dsDNA positivity* | 370 | 143 (38.6%) | 270 | 83 (30.7%) | 0.70 (0.51-0.98) | 0.04 |
| *Admission rate*<sup>2</sup> | 351 | 140 (39.9%) | 196 | 86 (43.9%) | 1.20 (0.84-1.71) | 0.31 |
| *Biologics*<sup>3</sup> | 332 | 23 (6.9%) | 239 | 12 (5.0%) | 0.70 (0.34-1.43) | 0.33 |

Summary of the available clinical history for the 656 study participants from cohort 3 since their disease diagnosis and up to the time of visit. Data were stratified by the study cohorts. <sup>1</sup>Odds ratio in non-European patients. *P* values were calculated using Pearson's chi-squared test. <sup>2</sup>Admission is defined as any patient requiring hospital admission specifically for their SLE in the 5 years prior to and including the date of their clinic visit at which the blood sample was taken. <sup>3</sup>Patients treated with Biologic drugs at any time since their disease diagnosis. EUR, patients of European ancestry; non-EUR, patients of non-European ancestry.

In addition to the ancestry-related changes, we found that sSIGLEC-1 levels were associated with lower levels of serum complement 3 (C3) (*P* = 5.0 × 10^−3^; Figure 4D), but not with other common serological markers of SLE such as C-reactive protein or anti-nuclear autoantibody levels (Supplemental Figure 2). This association was observed in both European and non-European patients and remained present even when adjusting for the presence of nephritis. The risk of renal complications/nephritis was higher in the combined population in patients with high concentrations of sSIGLEC-1 (OR = 1.53, *P* = 0.01; Table 2). These differences were much more pronounced in European patients (OR = 1.65, *P* = 0.02) compared to the non-European patients (OR=1.16; *P*=0.67; Table 2), and were maintained after adjusting for the association of sSIGLEC-1 with low serum complement C3 and C4 levels, which are predictors for renal disease (OR = 1.96 CI 1.10-3.45; *P* = 0.021). In addition, we also observed a similar trend towards increased risk of haematological complications, in patients with high concentrations of sSIGLEC-1, although not reaching statistical significance in our analysis (OR = 1.35, *P* = 0.09; Table 2).

**Table 2.** Association of sSIGLEC-1 with clinical parameters of SLE at the time of visit

|  |  |  |  |  |  |  |  |  |  |
| --- | --- | --- | --- | --- | --- | --- | --- | --- | --- |
|  | Combined |  |  | EUR |  |  | non-EUR |  |  |
| Clinical parameter | N (total) | OR<br>(95% CI) | p value | N | OR<br>(95% CI) | p value | N | OR<br>(95% CI) | p value |
| *Renal (nephritis)* | 156 | 1.53<br>(1.10-2.15) | 0.01 | 75 | 1.65<br>(1.09-2.52) | 0.02 | 81 | 1.16<br>(0.60-2.25) | 0.67 |
| *Haematological* | 291 | 1.35<br>(0.95-1.92) | 0.09 | 185 | 1.34<br>(0.80-2.20) | 0.25 | 106 | 1.28<br>(0.72-2.25) | 0.4 |
| *Biologics ever needed<sup>1</sup>* | 35 | 1.20<br>(1.01-1.42) | 0.04 | 23 | 1.18<br>(0.90-1.18) | 0.19 | 12 | 0.95<br>(0.70-1.23) | 0.71 |
| *Neurological* | 78 | 1.01<br>(0.76-1.34) | 0.95 | 36 | 0.87<br>(0.63-1.18) | 0.37 | 42 | 0.83<br>(0.39-1.76) | 0.64 |
| *Admission* | 226 | 1.18<br>(0.81-1.71) | 0.39 | 140 | 1.45<br>(0.90-2.37) | 0.13 | 86 | 0.60<br>(0.30-1.21) | 0.16 |
| *Corticosteroid use* | 302 | 0.98<br>(0.69-1.41) | 0.96 | 174 | 0.96<br>(0.57-1.57) | 0.86 | 128 | 0.88<br>(0.49-1.56) | 0.35 |
| Association of sSIGLEC-1 concentration and clinical parameters recorded for the SLE patients at the time of visit. Odds ratios were calculated for patients with sSIGLEC levels at >95th percentile. <sup>1</sup>Patients treated with Biologic drugs at any time since their disease diagnosis. OR, odds ratio; EUR, patients of European ancestry; non-EUR, patients of non-European ancestry. |  |  |  |  |  |  |  |  |  |

Further supporting the potential use of sSIGLEC-1 as a biomarker of disease severity, a higher frequency of patients with high levels of sSIGLEC-1 had a history of treatment with biologics (OR = 1.2, *P* = 0.04 in the combined population; Table 2), usually associated with patients with poor disease management that have not responded to standard treatment options. This remained the case even when we adjusted for history of renal complication as a confounding factor for biologics use.

## Discussion

Recent advances in medical research have led to the development of a breadth of novel treatment options. The characterisation of biomarkers that identify the exact pathophysiological mechanism underpinning the clinical manifestations in each patient has thus become a priority to allow the advent of a truly personalised medicine approach to human complex diseases. In systemic autoimmune and inflammatory diseases, chronic IFN signalling has been shown to be directly involved in the pathogenesis of the diseases, most notably in SLE(23). This observation has led to the development of therapeutic strategies to target IFN-α, which are currently being tested in the clinic (24). There is therefore an urgent need to develop robust and sensitive biomarkers that would allow the identification of patients with an active IFN response who would be more likely to benefit from anti-IFN-α therapy.

In the present study, we show that a circulating form of the surface-bound SIGLEC-1 receptor can be detected in human plasma/serum, and developed a sensitive immunoassay to measure the circulating concentrations of sSIGLEC-1. To our knowledge we are the first to detect the presence of a soluble SIGLEC-1 isoform in humans. Our current data do not allow us to determine the origin of sSIGLEC-1, and further work is needed to assess whether the soluble isoform is generated through alternative splicing or by proteolytic shedding of the membrane-bound receptor. A key feature of this bioassay is the limited sample requirements, making it amenable–as compared to a flow cytometric assay of surface SIGLEC-1– to screen large numbers of samples. We therefore propose that quantification of sSIGLEC-1 using our immunoassay is an alternative to flow cytometric endpoints, or the classical IFN signature, or its PCR-analysed surrogate(6), and will result in a more robust, simpler and less expensive measure of SIGLEC-1 expression and the IFN signature.

Other plasma/serum IFN-regulated biomarkers have been described in the literature, most notably IFN-α and IFN-γ-inducible protein 10 (IP-10) (12). However, the protein stability, cell-type specificity and much higher concentrations of sSIGLEC-1 are major practical advantage of our immunoassay. Moreover, measuring all 16 different IFN-α subtypes currently requires access to naturally-occurring high-affinity anti-IFN-α antibodies that are not readily available (22), which in combination with the much lower sample requirement compared to the SIMOA assay makes the sSIGLEC-1 assay ideally suited for high-throughput screening, including the retrospective testing of samples collected from completed clinical studies or cohorts for flares of IFN signalling and/or response to treatment of large cohorts and retrospective studies.

Clinically, we identified marked ancestry-related differences in sSIGLEC-1 levels among SLE patients, which were consistent with the higher disease severity in patients of non-European ancestry. In agreement with this hypothesis, sSIGLEC-1 levels were also associated with lower levels of C3, a classical serologic marker of the disease. Furthermore, high sSIGLEC-1 levels were found to predict renal and haematological complications, particularly in patients of European ancestry. These findings clearly underscore the importance of large, well-characterised, clinical cohorts to estimate the confounding effects of ancestry. A limitation of the current study was the lack of non-European healthy controls, which prevented us from investigating whether steady-state sSIGLEC-1 levels could also be altered in populations of non-European ancestry. However, the association of sSIGLEC-1 concentrations with overall increased disease severity was consistently maintained in both groups of patients. A possible explanation for the more modest predictive capacity observed in non-European patients is an increased disease heterogeneity, which could reflect a reduced dependency on the type I IFN pathway for disease severity and late-stage organ damage in this group of patients.

Taken together, our findings suggest that sSIGLEC-1 is a sensitive marker of monocyte and macrophage activation, which is critically implicated in the progression of several autoimmune and inflammatory diseases, such as SLE. In combination with additional available IFN-regulated biomarkers, the sSIGLEC-1 bioassay could help improve our capacity to dissect the molecular and clinical heterogeneity of complex conditions associated with an overt IFN response, and identify subsets of common and rare autoimmune and inflammatory diseases, collectively classified as interferonopathies. We have also shown that increased sSIGLEC-1 concentrations could, with further studies, have a clinical application in predicting increased risk of developing renal complications, one of the most severe clinical complication of SLE. Further work will now be required to validate these findings in additional autoimmune and inflammatory diseases associated with a chronic activation of the innate immune system using large and clinically well-characterised patient populations, as well as to extend the findings to non-European cohorts.

There has been considerable interest in developing drugs targeting the IFN-α signalling pathway to treat conditions associated with chronic IFN signalling. Our assay could also have utility in clinical trials, for example, for the selection of patients who could benefit the most from such inhibition of IFN-α. Conversely, IFN-α is also one of the most used compounds in cytokine therapy. However, its immunomodulatory properties may result in various autoimmune manifestations, with reported incidence of 4% to 19% in patients undergoing IFN-α therapy (25). We therefore hypothesize that these adverse events may be avoided if the background IFN signature is known and therapy adjusted to avoid the excessive IFN signalling known to be a factor in the promotion of secondary autoimmunity in these patients. Moreover, it has been recently suggested that the detection of an IFN signature in peripheral blood is associated with poor response to both B-cell depleting therapy (rituximab) and anti-IL-6R treatment (tocilizumab) in rheumatoid arthritis patients (26, 27). These data suggest that sSIGLEC-1 could be useful in prediction to therapeutic responses, and supports a broader application of this assay in the context of patient stratification for clinical trials.

## Methods

### Subjects

Discovery cohort (cohort 1) study participants included 34 SLE patients (median age, 39 years, range 20-72, 32/34 female) recruited from Guy’s and St Thomas’ NHS Foundation Trust. All patients satisfied ACR SLE classification criteria and were allocated a disease activity using SLEDAI-2K at the time of sampling. Patients were recruited from a clinic in which the severity of disease was such that none of the patients was on high dose oral corticosteroids (> 15 mg/day) or B-cell depleting therapy. Healthy volunteers matched for age and sex (n=24, median age, 43, range 25-60 years 23/24 female) were recruited from the Cambridge BioResource.

A replication cohort (cohort 2) of 41 SLE patients (median age, 52 years, range 21-82, 38/41 female) and 490 healthy volunteers (median age, 48 years, range 18-78, 320/490 female) was recruited from the Cambridge BioResource. The SLE patients were recruited specifically for this study outside their regular clinic visits, and were otherwise well at the time of bleeding. No additional disease information or ancestry data was available for this cohort of patients.

To investigate the association between sSIGLEC-1 with ancestry and clinical manifestations, a third independent cohort (cohort 3) of SLE patients (n=656, median age, 45 range 15-82 years 592/655 female, one unknown) was recruited from multiple collaborative centres in the UK (St Thomas’s Hospital, Newcastle Hospital, City Hospital Sunderland, City Hospital Birmingham, Royal Hallamshire Hospital, Hammersmith Hospital, West Middlesex Hospital and Basildon Hospital). Patients were of European (n=370), South East Asian (n=134), African/Afro-Caribbean (n=94) and Far East Asian (n=19) ancestries. Twenty-three patients were of other minor ancestry groups (including Middle Eastern, Maori and Fiji ancestry) and 16 had missing ancestry information. The history of clinical manifestations since their disease diagnostic up to patients is summarised in Table 1.

All samples and information were collected with written and signed informed consent after approval from the relevant research ethics committees (refs: 05/Q0106/20, 07/H0718/49 and 08/H0308/153).

### Flow cytometry

SIGLEC-1 expression was measured in PBMCs from 34 SLE patients and 24 healthy donors from the discovery cohort. PBMCs were thawed in a 37°C waterbath, resuspended in X-VIVO 15 (Lonza) + 1% heat-inactivated, filtered, human AB serum (Sigma) and immunostained for 30 min at room temperature. SIGLEC-1 expression on CD14^+^ monocytes was determined using fluorochrome-conjugated antibodies against CD14 (Clone M5E2, BioLegend) and SIGLEC-1 (Clone 7-239, BioLegend). Immunostained samples were acquired using a BD LSR Fortessa (BD Biosciences) flow cytometer, and data were analysed using FlowJo (Tree Star). Dead cells were excluded based on the eFluor780 Fixable Viability Dye (eBiosciences).

### Soluble SIGLEC-1 time-resolved fluorescence immunoassay

Plasma/serum sSIGLEC-1 concentrations were measured using a non-isotopic time-resolved fluorescence (TRF) assay based on the dissociation-enhanced lanthanide fluorescent immunoassay technology (DELFIA; PerkinElmer). Duplicate test plasma/serum samples diluted 1:10 in assay buffer (PBS, 0.05% Tween-20, 10% FCS) were incubated 2 h at room temperature and then at 4°C overnight in 96-well EIA/RIA plates (Corning) coated with 1 μg/ml mouse monoclonal anti-human SIGLEC-1 capture antibody (AB18619; Abcam). Sample detection was performed using a biotinylated sheep polyclonal detection antibody (BAF5197; R&D Systems) diluted to a final concentration of 200 ng/ml. Following incubation with the secondary antibody, europium-labelled streptavidin (PerkinElmer) was added and concentration of antigen was measured by the amount of disassociated europium that is fluorescent at 615 nm after excitation at 320 nm.

Quantification of test samples was obtained by fitting the readings to a human recombinant SIGLEC-1 (R&D Systems) serial dilution standard curve on each plate (r^2^>0.995). To maintain assay consistency, the recombinant protein was aliquoted and stored at −80°C immediately following reconstitution and a fresh aliquot from the same lot was used for each assay.

The lower limit of detection of the assay was set as 2- background levels in each plate and corresponded to an average of 1.29 ng/ml across all plates. Samples with measurements below the limit of detection (6/589) were set to 1.29 ng/ml. Assay specificity was confirmed using a biotinylated sheep polyclonal isotype control (R&D Systems). Technical variation was assessed in duplicate measurements of all samples (average CV = 5.0%). Samples with CV > 30% between duplicates (10/589) were excluded from the analysis. To evaluate potential matrix effects, we diluted a sample with high sSIGLEC-1 concentration with assay medium and showed a linear titration between 1:2 and 1:16 dilutions (r^2^ = 0.94).

### IFN-α_2b_ single-molecule digital ELISA assay

Circulating levels of IFN-α_2b_ were measured by Single-Molecule Array (SIMOA) digital ELISA (Quanterix) according to the manufacturer’s instructions. IFN-α detection was achieved using mouse anti-IFN-α monoclonal antibodies: a neutralising antibody (clone MMHA – capture) and an anti-IFN-α antibody raised against an IFN-α_2b_ antigen (clone 7N41 – detection); cross-reactivity to the other IFN-α subtypes was not assessed. Measurements were performed in plasma samples that had never been previously thawed from the 41 SLE patients from cohort 2, which had never been previously thawed.

### Type I IFN transcriptional signature in PBMCs

RNA from 34 SLE patients and 24 age- and sex-matched healthy donors in cohort 1 and from 41 SLE patients and 41 age- and sex-matched healthy donors in cohort 2 was extracted from freshly isolated PBMCs stored in TRIZOL immediately after collection, using the Direct-zol RNA Mini-Prep kit (Zymo Research), following manufacturer’s instructions. RNA concentration was measured by NanoDrop (Thermo Fisher Scientific), and 50 ng of total RNA were hybridized with a custom NanoString CodeSet (NanoString Technologies), containing 56 IRGs previously identified as discriminative of the IFN signature (21). Expression levels were assessed using an nCounter Flex instrument (NanoString Technologies). Data were processed using the nSolver Analysis Software following normalization to the geometric mean of positive control spike-ins, and the gene expression of eight housekeeping genes.

Expression of 56 IRGs previously identified as discriminative of the IFN signature(21), were assessed using RNA from: (i) 34 SLE patients and 24 age- and sex-matched healthy donors in cohort 1; and (ii) 41 SLE patients and 41 age- and sex-matched healthy donors in cohort 2, using a custom NanoString CodeSet (NanoString Technologies).

A quantitative metric of the IFN signature was generated using principal component analysis by the projection of the expression of the 56 IRGs onto the first principal component (PC1), which was found to explain 86.3% of the variance of this dataset. A complete list of the 56 IRGs and respective NanoString probe sequences are listed in Supplemental Table 1.

### Statistical analyses

Statistical analyses were performed using Prism software (GraphPad) and R (www.r-project.org). Given that most phenotypes showed moderate to strong right skew that violated the assumption of normality, the phenotypes were log-transformed before statistical testing and all reported values refer to the geometric mean of the respective measurements.

*Cohorts 1 and 2*: the association of sSIGLEC-1 and measured immune parameters in cohorts 1 and 2 was performed using two-tailed non-parametric Mann-Whitney tests. Correlation analyses were performed using linear regression on the log-transformed data.

*Cohort 3*: all statistical analyses with the clinical data available for the 656 patients from cohort 3 were performed using R software. Association between ancestry and sSIGLEC-1 concentration was assessed using a paired student t-test. Odds ratio (OR) of each clinical parameter (history of admission within 5 years, *ever* having suffered with renal, haematological or neurological disease, requiring biological therapy or active corticosteroid use to control disease) in European and non-European patients was assessed by Pearson’s Chi-squared test.

Patients were divided into groups based on sSIGLEC-1 serum level centiles (< 50^th^ centile, 51-74^th^ centile, 75-95^th^ centile and >95^th^ centile). Association of each group and clinical parameters was performed using a logistical regression model. Patients of European and non-European ancestry were analysed separately as ethnicity was a major confounding factor.

Association of sSIGLEC-1 concentration and other serological parameters of disease (complement levels (C3 and C4 – measured within 3 months of the visit at the patient’s local centre/hospital), anti-dsDNA antibody titres and C-reactive protein) were assessed using linear regression on the log transformed data.

## Acknowledgements

We gratefully acknowledge the participation of all NIHR Cambridge BioResource volunteers and thank the NIHR Cambridge BioResource centre and staff for their contribution. We thank the National Institute for Health Research and NHS Blood and Transplant. We thank Neil Walker and Helen Schuilenburg from the Cambridge Institute for Medical Research, University of Cambridge for data management. We thank members of the Cambridge BioResource SAB and management committee for their support and the National Institute for Health Research Cambridge Biomedical Research Centre for funding. We also thank Helen Stevens, Gill Coleman, Sarah Dawson, Simon Duley, Meeta Maisuria and Sumiyya Mahmood from the Cambridge Institute for Medical Research, University of Cambridge for preparation of DNA and PBMC samples. We also thank Emma Jones from AstraZeneca for the use of the NanoString instrument and Claudia Gonzalez-Lopez from the Weatherall Institute of Molecular Medicine, Oxford, for the use of the Simoa HD-1 analyzer.

---

**Supplemental Figure 1.**
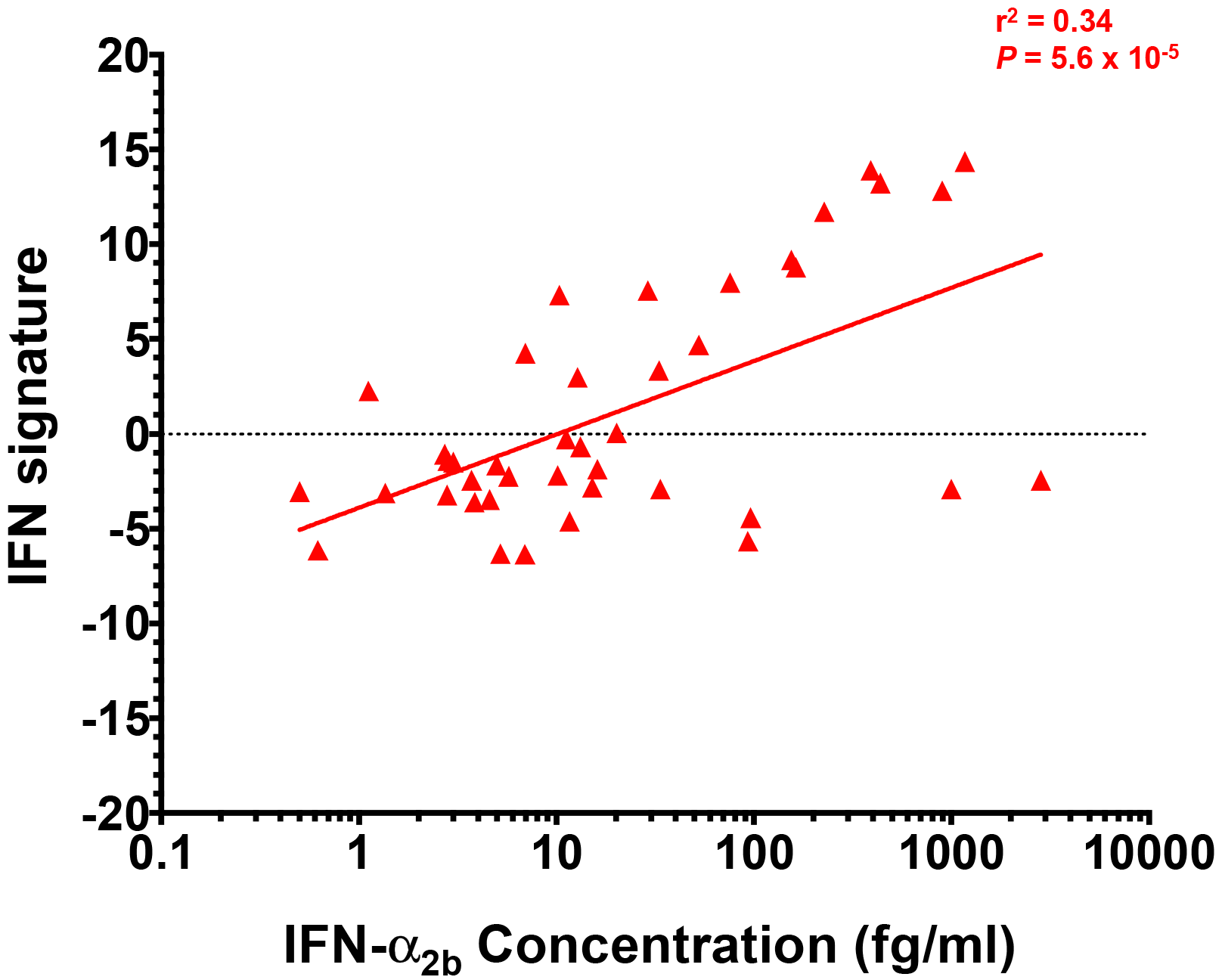
Concentration of IFN-α_2b_ is associated with the IFN transcriptional signature. Data shown depict the correlation between the plasma IFN-α_2b_ concentration and the transcriptional IFN signature in 41 SLE patients in cohort 2.

**Supplemental Figure 2.**
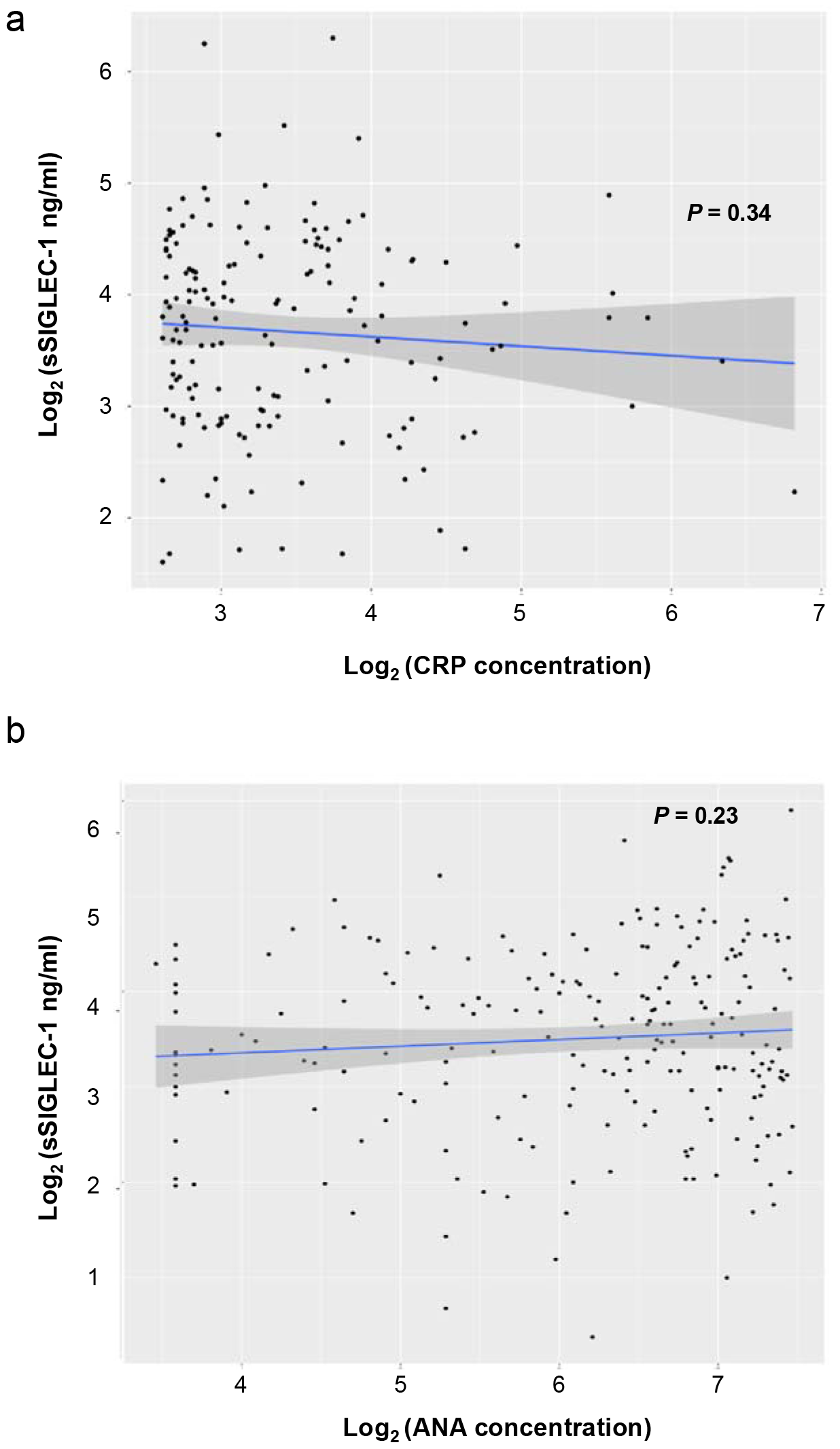
Association of sSIGLEC-1 with serological markers of SLE. (A,B) Data shown depict the association of sSIGLEC-1 concentrations with C-reactive protein (CRP) levels (A) and with disease-specific anti-nuclear autoantibody (ANA) titres (B). *P* values were calculated by linear regression.

